# Structural insight into the substrate binding of the AMT complex via an inhibitor-trapped state

**DOI:** 10.1101/2025.04.24.650485

**Authors:** Zengyu Shao, Sol Yoon, Jiuwei Lu, Pranav Athavale, Yifan Liu, Jikui Song

**Affiliations:** Department of Biochemistry, University of California, Riverside, CA 92521, USA; Department of Cancer Biology, University of Southern California, Los Angeles, CA 90033, USA

## Abstract

N6-adenine (6mA) DNA methylation plays an important role in gene regulation and genome stability. The 6mA methylation in *Tetrahymena thermophila* is mainly mediated by the AMT complex, comprised of the AMT1, AMT7, AMTP1 and AMTP2 subunits. To date, how this complex assembles on the DNA substrate remains elusive. Here we report the structure of the AMT complex bound to the OCR protein from bacteriophage T7, mimicking the AMT-DNA encounter complex. The AMT1-AMT7 heterodimer approaches OCR from one side, while AMTP1 N-terminal domain, assuming a homeodomain fold, binds to OCR from the other side, resulting in a saddle-shaped architecture reminiscent of what was observed for prokaryotic 6mA writers. Mutation of the AMT1, AMT7 and AMTP1 residues on the OCR-contact points led to impaired DNA methylation activity to various extents, supporting a role for these residues in DNA binding. Furthermore, structural comparison of the AMT1-AMT7 subunits with the evolutionarily related METTL3-METTL14 and AMT1-AMT6 complexes reveal sequence conservation and divergence on the region corresponding to the OCR-binding site, shedding light onto the substrate binding of the latter two complexes. Together, this study suggests a substrate binding-induced open-to-closed conformational transition of the AMT complex, with implications in its substrate binding and processive 6mA methylation.

## Introduction

DNA methylation is an epigenetic mechanism critically influencing chromatin structure and function. The most well-known DNA methylation occurs at the C-5 position of cytosines, which functions to regulate gene expression^1,2^, X-chromosome inactivation^3^, transposon silencing^4,5^, and genome stability^6^. In addition, DNA methylation at N6-adenine (6mA) plays a role in modulating genome function in both prokaryotes and eukaryotes^7,8^. Recent studies have further linked 6mA DNA methylation in protists, green algae and basal fungi, occurring predominantly in the context of ApT dinucleotides, to epigenetic regulation due to its association with transcriptionally active domains^9-12^ and epigenetic marks H2A.Z and H3K4me3^13,14^. However, in contrast to the writers of C-5 DNA methylation that have been well characterized^15-20^, the structure and mechanism of the writers for 6mA DNA methylation in eukaryotes remains elusive.

In unicellular eukaryote *Tetrahymena thermophila*, 6mA DNA methylation is mainly maintained by the highly processive AMT complex (a.k.a. MTA1c) comprised of AMT1 methyltransferase (MTase) (a.k.a. MTA1), and regulatory subunits AMT7 (a.k.a. MTA9), AMTP1 (a.k.a. p1), and AMTP2 (a.k.a. p2)^9,12^. In addition, a variant of the AMT complex, in which AMT7 is replaced by a shorter homologue AMT6, plays a role in maintenance of 6mA methylation in a replication-coupled manner^14^. Previous structural and biochemical characterizations of the AMT complex from *Tetrahymena thermophila* reveal that AMT1, AMT7, AMTP1, and AMTP2 form a heterotetramer in a stoichiometry of 1:1:1:1^21,22^. Among these, the catalytic AMT1 subunit harbors a classic DPPW catalytic motif, stacked with the regulatory AMT7 subunit in a manner resembling the heterodimeric RNA adenine methyltransferase METTL3-METTL14 complex^23-25^. Furthermore, a long N-terminal helix of AMT1 bridges the AMT1-AMT7 heterodimer with the AMTP1 and AMTP2 subunits through interactions with AMTP2 and C-terminal helix (CH) of AMTP1. It has been established that the AMT complex-mediated 6mA methylation requires the presence of all four subunits, with the homeodomain-like DNA-binding proteins AMTP1 and AMTP2 critically regulating substrate binding^21,22^. However, due to lack of structural information on substrate recognition, the molecular mechanism by which four members of AMT complex orchestrate 6mA methylation remains elusive.

To elucidate the mechanism of the substrate binding by the AMT complex, we resorted to the Overcoming Classical Restriction (OCR) protein from bacteriophage T7, a mimic of B form DNA^26^, to stabilize the potential DNA-binding surfaces of AMT1, AMT7, AMTP1, and AMTP2. The OCR protein binds to the AMT complex tightly, leading to inhibition of the latter’s DNA methylation activity. On this basis, we solved the single-particle cryo-EM structure of AMT1-AMT7-AMTP1-AMTP2-OCR complex. The cryo-EM structure of the AMT complex reveals saddle-shaped architecture, in which the AMT1-AMT7 heterodimer is bridged with the AMTP1 and AMTP2 subunits via the AMT1 N-terminal helix (NH). Strikingly, the AMT1-AMT7 subunits are joined by the N-terminal domain of AMTP1, which is otherwise structurally disordered in the apo form of the AMT complex, to flank the conformationally dynamic OCR, resulting in a closed conformation reminiscent of what was observed for many other 6mA writers in prokaryotes. Our combined structural and biochemical analysis confirmed the OCR-binding surface as potential DNA-binding sites and shed light onto the substrate-binding mechanism of the evolutionarily related METTL3-METTL14 complex. Together, this study supports the notion that the AMT complex undergoes an open-to-closed conformational transition upon substrate binding, with important implications in the mechanism of the AMT complex-mediated 6mA DNA methylation.

## Results

### The OCR protein inhibits the AMT complex-mediated 6mA DNA methylation

A previous structural study of the AMT complex, despite in the presence of DNA substrate, failed to observe the density of the DNA^22^, suggesting that the AMT-DNA association is highly dynamic. To overcome this challenge, we resorted to the OCR protein from T7 bacteriophage for stabilizing the substrate-binding conformation of the AMT complex (Fig. 1a-c). OCR reportedly exists in a homodimeric form mimicking a slightly bent 20-base pair B-form DNA and has been shown to bind and inhibit DNA-templated enzymes^27-29^. To test the interaction between the AMT complex and OCR, we performed size-exclusion chromatography analysis for the AMT complex sample mixed with OCR. OCR co-migrated with the AMT complex on a Superdex 200 increase 10/300 GL column, indicative of stable association (Fig. 1d, e). To evaluate whether the presence of OCR affects the AMT complex-mediated 6mA DNA methylation, we performed in vitro DNA methylation assay for the AMT complex on a 12-mer DNA duplex containing a single hemimethylated ApT site in the presence or absence of OCR. Consistent with previous observations^9,12,13,21,22^, the AMT complex is highly active on the hemimethylated ApT DNA (Fig. 1f). However, the presence of OCR in a 2:1 molar excess reduced the DNA methylation efficiency of the AMT complex by ∼100-fold (Fig. 1f), indicative of strong inhibition. These data establish OCR as a DNA-mimicking inhibitor for the AMT complex-mediated 6mA DNA methylation.

**Fig. 1:**
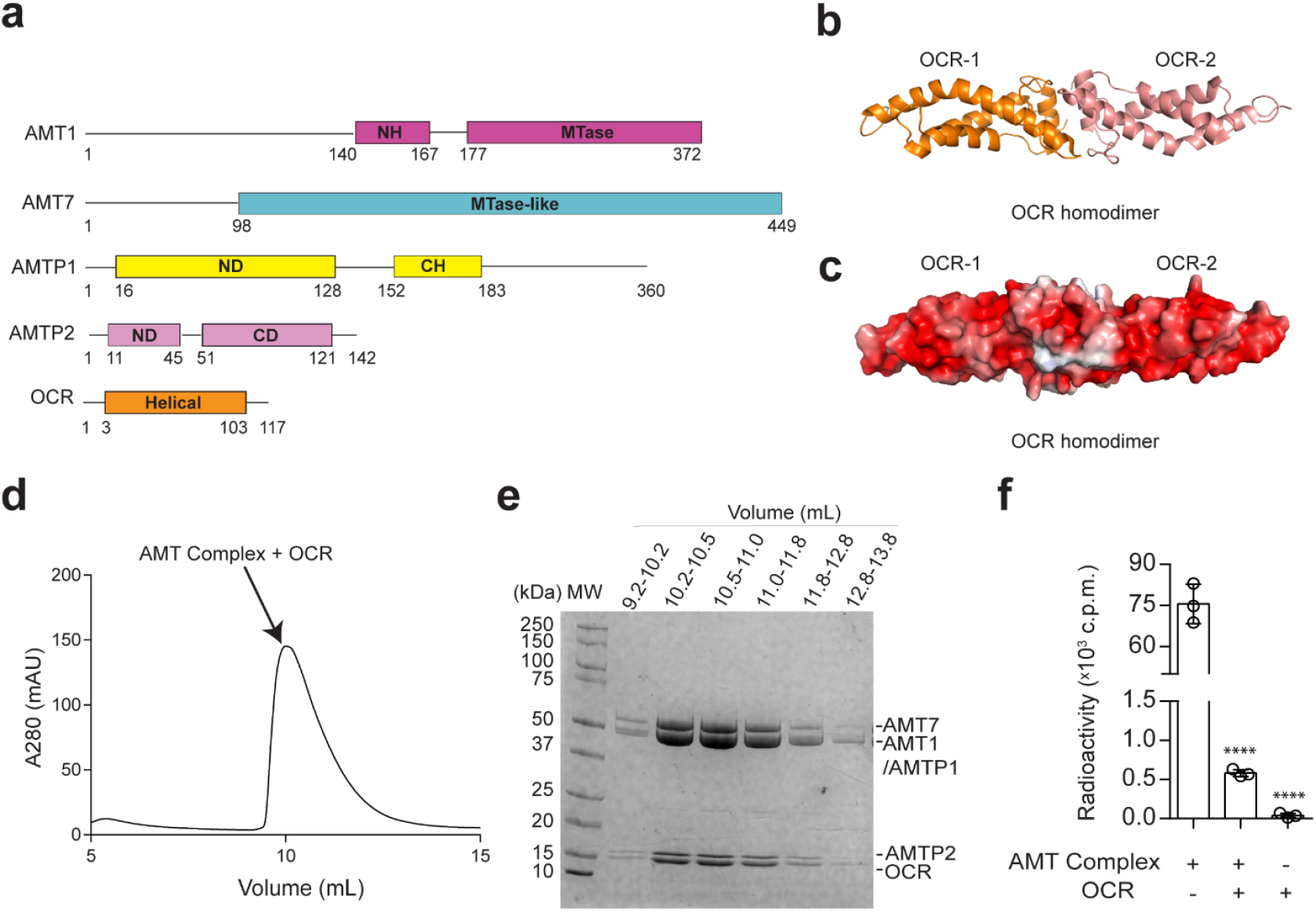
Biochemical analysis of the AMT-OCR complex. **a** Domain architecture of AMT1, AMT7, AMTP1, AMTP2, and OCR. NH, N-terminal helix; ND, N-terminal domain; CH, C-terminal helix; CD, C-terminal domain. **b, c** Ribbon (b) and surface electrostatic (c) representation of OCR homodimer (PDB 8ZEK). **d, e** Size-exclusion chromatography (d) and the corresponding SDS-PGAE analysis of selected fractions (e) of the AMT-OCR sample mixture. **f** In vitro DNA methylation assay of the AMT complex on a hemimethylated 12-mer DNA duplex, in the presence or absence of OCR. Data are mean ± s.d. (n = 3 biological replicates). The two-tailed Student’s t test statistical analysis was performed to compare AMT complex activity in the absence vs. presence of OCR. ****p < 0.0001.

### Structural overview of the OCR-bound AMT complex

Next, we solved the cryo-EM structure of the OCR-bound AMT complex at an overall 3.59-Å resolution (Fig. 2a, b, Supplementary Fig. 1 and Supplementary Table 1). Structural analysis of the OCR-bound AMT complex reveals that the AMT complex is comprised of one copy of AMT1, AMT7, AMTP1, and AMTP2 each (Fig. 2a-d and Supplementary Fig. 2), consistent with previous observations^21,22^. We were able to trace the density for AMT1 residues 128-216 and 227-372, AMT7 residues 99-114, 118-156, 162-228, 319-450, AMTP1 residues 16-128 and 154-184, and AMTP2 residues 11-132 (Fig. 2c, d and Supplementary Fig. 2).

**Fig. 2:**
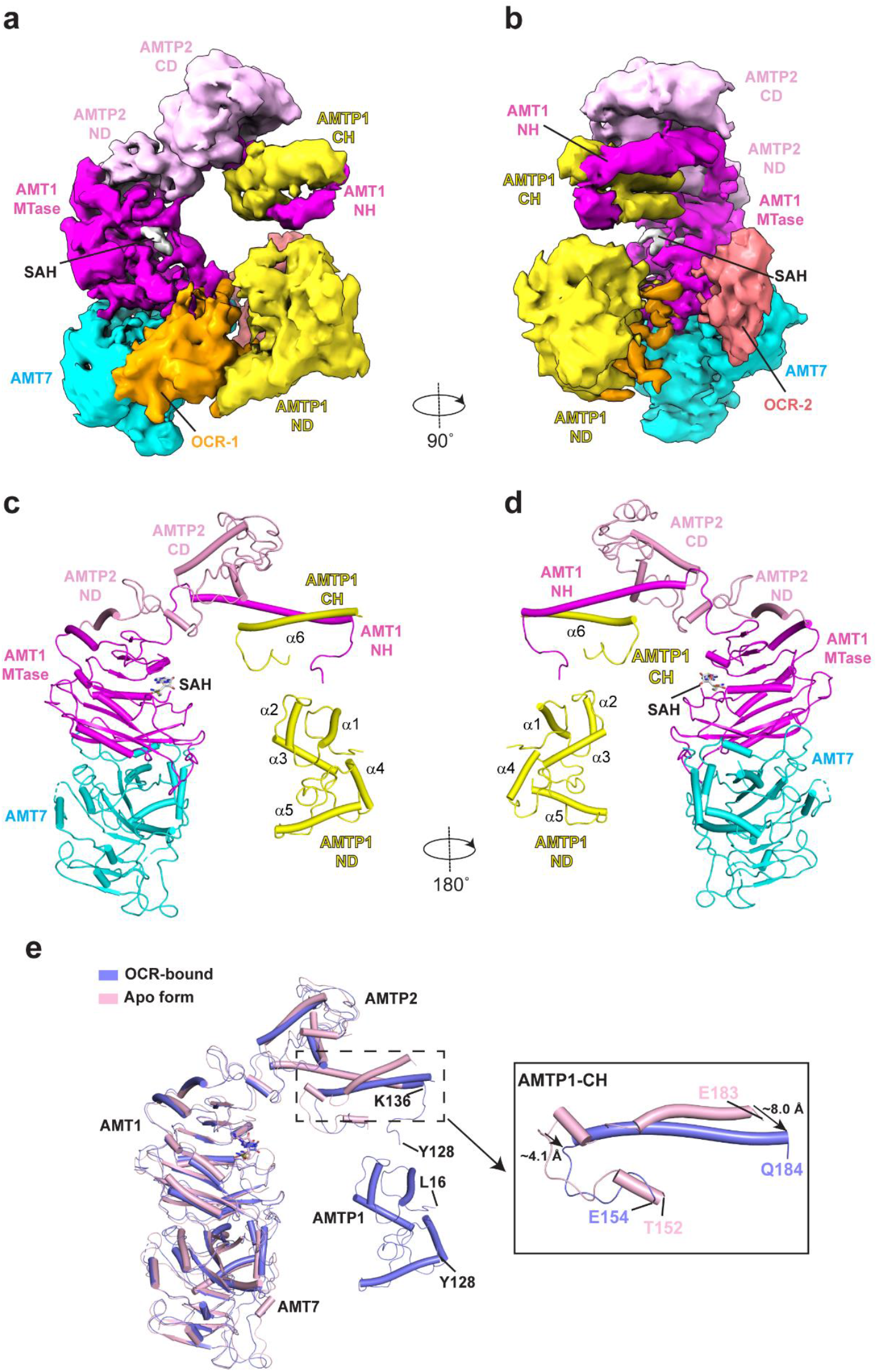
Cryo-EM structure of the OCR-bound AMT complex. **a, b** Two orthogonal views of the cryo-EM density map of the AMT-OCR complex, with individual subunits and domains labeled. AMT1, AMT7, AMTP1, and AMTP2 are colored in magenta, cyan, yellow, and pink, respectively. The color scheme applies to subsequent figures unless otherwise indicated. The density corresponding to the OCR subunits (OCR-1 and OCR-2) are colored in orange and deep salmon, respectively. **c, d** Atomic model of the AMT-OCR complex in two opposite views. The cofactor by-product SAH molecule is shown in stick representation. **e** Structural overlay of apo-form (PDB 7YI8) and OCR-bound AMT complex, with the conformational shift for the AMTP1 CH highlighted in the expanded view.

The density for the OCR molecules bound to the AMT complex is rather weak, presumably due to conformational flexibility, which prevents us from accurate modeling (Fig. 2a, b and Supplementary Fig. 2). Nevertheless, it is apparent that the OCR molecules are sandwiched by the N-terminal domain (ND) of AMTP1 from one side and the AMT1 MTase and AMT7 MTase-like domains on the other side (Fig. 2a, b). As observed previously^21,22^, the MTase domain of AMT1 associates with the MTase-like domain of AMT7 in antiparallel, resulting in a butterfly-like fold (Fig. 2c, d). The AMTP1 protein is comprised of an N-terminal domain (ND) dominated by a helical fold and a structurally independent C-terminal helix (CH) (Fig. 2c, d). The AMTP2 protein is comprised of an ND in a helix-loop-helix conformation and a C-terminal domain (CD) dominated by a four-helix bundle (Fig. 2a-d). Notably, the AMT1 N-terminal helix (NH) engages in helical contacts with both the AMTP1 CH and the AMTP2 CD, thereby bridging the AMT1-AMT7 heterodimer with AMTP1 and AMTP2 (Fig. 2a-d). Meanwhile, the ND of AMTP2 is docked onto the surface region formed by the beginning α-helix and β-strand of the AMT1 MTase domain as well as the preceding loop (Fig. 2c-d).

Structural superposition of the OCR-bound AMT complex and the previously reported apo-form AMT complex (PDB 7YI8)^22^ gave a root-mean-square deviation (RMSD) of 1.2 Å over 531 aligned Cα atoms, indicative of high structural similarity (Fig. 2e). The most notable difference between the two complexes lies in the AMTP1 ND domain, which is disordered in the apo-AMT complex but is well defined in the OCR-bound form (Fig. 2e), suggesting an OCR binding-induced structural stabilization effect. The conformational difference between the apo- and OCR-bound AMT complex was further propagated to the AMTP1 CH (residues E154-Q184) (Fig. 2e): upon the OCR binding, the N- and C-termini of the AMPT1 CH shifted toward the AMTP1 ND by ∼4 and ∼8 Å, respectively (Fig. 2e).

### Mapping of the potential DNA-binding interface of the AMT1-AMT7 heterodimer

Analysis of surface electrostatic potential of the OCR-bound AMT complex reveals that the OCR molecules are walled by the positively charged surfaces of the AMT1 MTase domain, the AMT7 MTase-like domain, and the AMTP1 ND domain (Fig. 3a-c). On one side, a cluster of residues at the AMT1-AMT7 interface, including AMT1 K280, K289 and H291 and AMT7 R379 and R382, form a contact surface with the OCR (Fig. 3b). Proximal to this surface are the residues (Y227, R357 and R358) around the cofactor *S*-Adenosyl methionine (SAM)-binding site of AMT1, formed by two loop segments, namely Gate loops 1 and 2^21,22^ (Fig. 3b). On the other side, AMTP1 residues K44, K46, K57, R68, K101, K114 and K118 form another contact surface for the OCR (Fig. 3c).

**Fig. 3:**
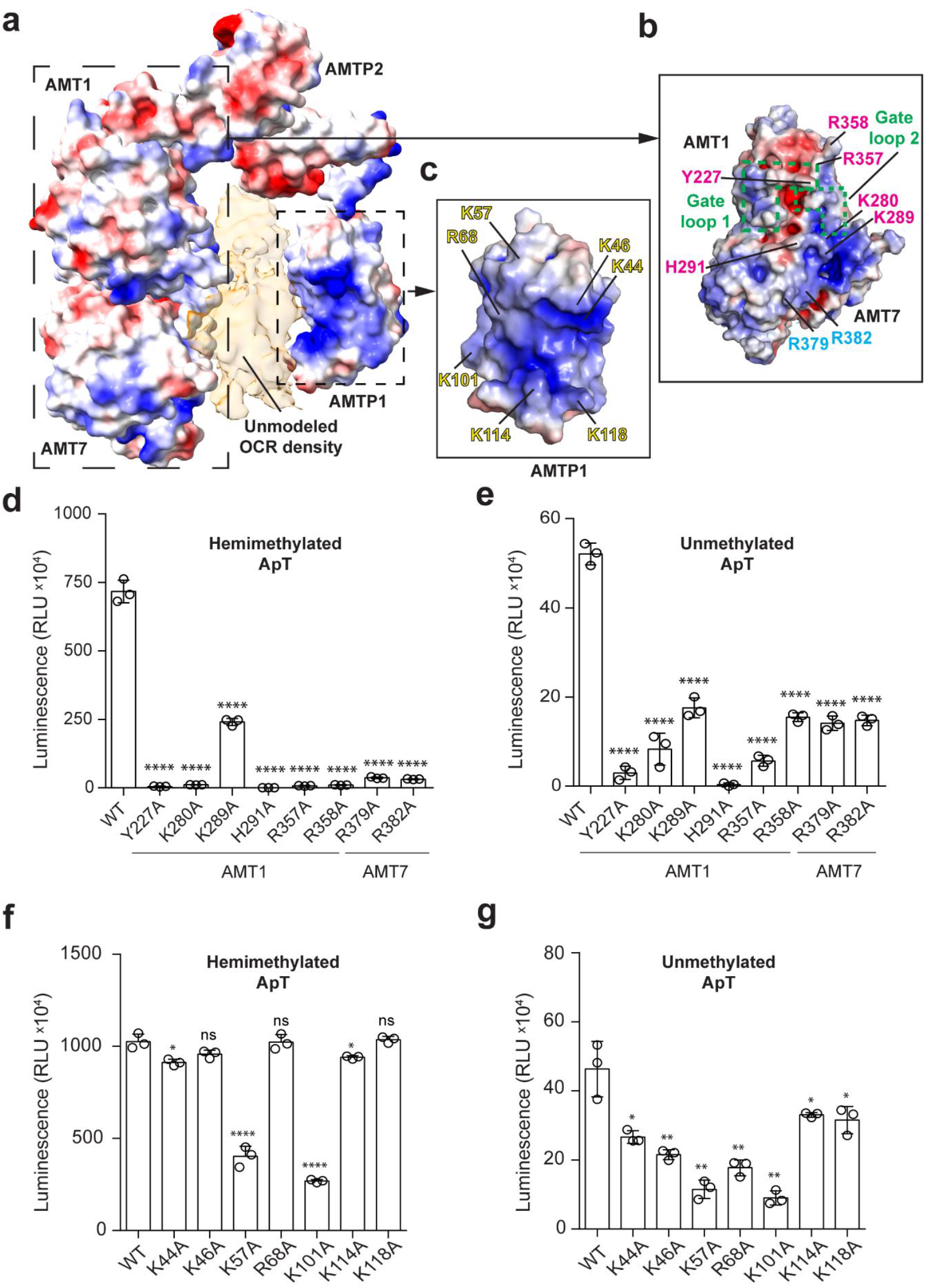
Structural and mutational analysis of the potential DNA-binding surface of the AMT complex. **a-c** Electrostatic surface of the AMT complex, with the potential substrate-binding sites of the AMT1-AMT7 dimer (b) and AMTP1 (c) shown in the expanded views. Gate loops 1 and 2 in AMT1 are highlighted by green dashed lines. **d, e** In vitro DNA methylation assay of the AMT1- or AMT7-mutated AMT complex on a hemimethylated 12-mer DNA duplex (d) or unmethylated 24-mer DNA duplex (e). Data are mean ± s.d. (n = 3 biological replicates). **f, g** In vitro DNA methylation assay of the AMTP1-mutated AMT complex on a hemimethylated 12-mer DNA duplex (f) or unmethylated 24-mer DNA duplex (g). Data are mean ± s.d. (n = 3 biological replicates). The two-tailed Student’s t test statistical analysis was performed to compare the activity of WT vs. mutant AMT complex. ns, not significant; *p<0.05, **p<0.01; ****p < 0.0001.

The fact that the OCR protein serves as a mimic for the B-form DNA suggests that the OCR-contact sites of the AMT complex constitute its DNA-binding interface. To test this possibility, we selected the OCR-contact residues of AMT1, AMT7, and AMTP1 for mutagenesis and performed in vitro DNA methylation assays on substrates containing either a favorable hemimethylated ApT site or unfavorable, unmethylated ApT sites (Fig. 3d-g). For the hemimethylated substrate, introducing the AMT1 K289 mutation led to ∼3-fold reduction of the DNA methylation activity, while introducing the AMT1 Y227A, K280A, H291A, R357A or R358A, or the AMT7 R379 or R382 mutation, largely abolished the AMT activity (Fig. 3d). Likewise, introducing the AMT1 and AMT7 mutations also led to substantial reduction of the AMT activity on the unmethylated substrate (Fig. 3e). For AMTP1, introducing the K57A and K101A mutations led to 2-4-fold lower methylation efficiency of the AMT complex on the hemimethylated substrate, and 3-6-fold lower methylation efficiency on the unmethylated substrate, suggesting an important role for these two sites in DNA binding (Fig. 3f, g). On the other hand, a less pronounced effect was observed for the AMTP1 K44A, K46A, R68A, K114A or K118A mutation, which led to slight or no activity change of the AMT complex on the hemimethylated substrate (Fig. 3f). Nevertheless, it is apparent that these mutations significantly impair the AMT-mediated methylation on the less favorable, unmethylated substrates, supporting a role for these residues in fine-tuning the activity of the AMT complex (Fig. 3g). Sequence analysis of AMT1, AMT7 and AMTP1 across various species further demonstrated that many of these potential DNA-binding sites, such as AMT1 Y227, K280, K289, H291, R357 and R358, AMT7 R379 and R382, and AMTP1 K44, K46, K57, R68 and K101, are evolutionarily conserved (Supplementary Fig. 3 and 4). Together, these data support the notion that the OCR-contact residues provide a DNA-binding interface underpinning the AMT-mediated 6mA DNA methylation.

### Comparative structural analysis of AMTP1 in AMT-mediated 6mA methylation

AMTP1 belongs to a broad family of homeodomains, which often interact with cognate DNAs to regulate cell- and tissue-specific gene expression^30^. Through structural homology search using the DALI program^31^, we identified a few structural homologues for AMTP1, including Double homeobox protein 4 (DUX4) (PDB 6E8C)^32^, Myb-containing domain (MCD3) of Double-strand telomeric DNA-binding proteins 2 (TEBP-2) (PDB 8U8L)^33^, Antennapedia homeodomain-DNA complex (PDB 9ANT)^34^, and TEA domain family transcription factor 4 (TEAD4) (PDB 5GZB)^35^ (Supplementary Fig. 5a). Notably, these proteins all harbor a homeodomain, employing a helix-turn-helix (HTH) motif to access the major groove of the DNA molecules (Supplementary Fig. 5b). It’s worth noting that several OCR-contact sites for AMTP1 are also located on the corresponding region, spanning from α2-helix to α3-helix of the ND, such as K44, K46, K57 and R68, in addition to those from the neighboring β1-strand (K101) and α5-helix (K114 and K118) (Fig. 2c and Supplementary Fig. 4 and 5c). These data provide further insight into the DNA-binding sites of AMTP1.

### Structural comparison of the AMT complex with prokaryotic 6mA writers

To illustrate how the AMT complex-substrate binding is linked to prokaryotic adenine DNA methyltransferases, we generated a structural model of the AMT-DNA complex, based on the AMT-OCR interaction, and performed structural comparison with *E. coli* Type III restriction-modification enzyme EcoP15I-DNA complex (PDB 4ZCF)^38^, *Caulobacter crescentus* cell cycle-regulated DNA methyltransferase (CcrM)-DNA complex (PDB 6PBD)^37^, and *Thermus aquaticus* DNA adenine methyltransferase M.TaqI-DNA complex (PDB 1G38)^36^(Fig. 4a-h). Similar to the AMT complex, the EcoP15I complex employs multiple subunits for DNA binding, with the catalytic ModB subunit binding to the DNA from the minor groove and the ModA subunit binding to the major groove for sequence-specific recognition (Fig. 4c,d). Likewise, the CcrM complex involves two copies of CcrM molecules for DNA binding, with one molecule approaching the minor groove for catalysis and the other approaching the major groove (Fig. 4e,f). In contrast, the M.TaqI complex relies on two subdomains of M.TaqI: catalytic core and the target recognition domain (TRD), to embed DNA from minor groove and major groove, respectively (Fig. 4g,h). Despite distinct assembly mechanisms, these complexes all form a deep cleft for processive DNA methylation (Fig. 4a-h), highlighting similar yet distinct mechanisms of substrate DNA binding among the evolutionarily related 6mA DNA methyltransferases.

**Fig. 4:**
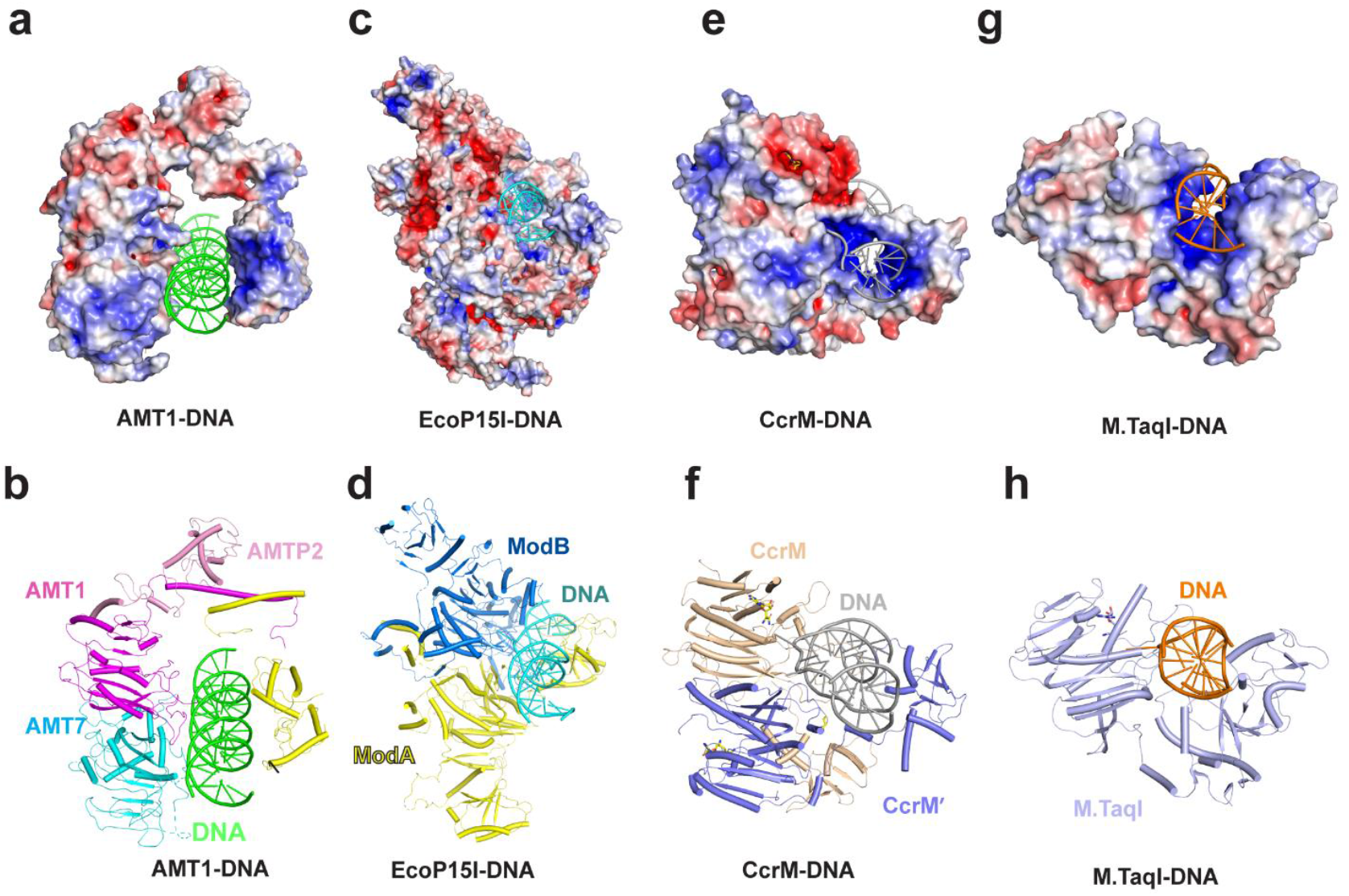
Structural comparison of the AMT-DNA model with the enzyme-substrate complexes of bacteria 6mA DNA methyltransferases. **a, b** Electrostatic surface (a) and ribbon representation (b) of the AMT-DNA model. **c, d** Electrostatic surface (c) and ribbon representation (d) of the M.TaqI-DNA complex (PDB 1G38). **e, f** Electrostatic surface (e) and ribbon representation (f) of the CcrM-DNA complex (PDB 6PBD). **g, h** Electrostatic surface (g) and ribbon representation (h) of the EcoP15I Mod subunit-DNA (PDB 4ZCF).

Structural comparison of the AMT complex with the prokaryotic 6mA writers further indicated that AMTP1 structurally corresponds to the major groove-binding components of the prokaryotic methyltransferase complexes, such as the TRD of M.TaqI, the TRD of the ModA subunit of EcoP15I and the DNA binding domain of the major grove-engaging CcrM molecule (Fig. 4a-h). Whether AMTP1 recognizes the target DNA in a similar manner warrants further investigation.

### Structural insights into the substrate-binding site of the METTL3-METTL14 complex

The AMT1-AMT7 heterodimer and the RNA adenine methyltransferase METTL3-METTL14 heterodimer belong to the same MT-A70 methyltransferase family^9,12^, with high sequence and structural similarity^21,22,24,25^. Structural overlay of the OCR-bound AMT complex with the METTL3-METTL14 complex (PDB 5IL1)^25^ reveals high structural similarity between the two complexes, with an RMSD of 1.13 Å over 161 aligned Cα atoms (Supplementary Fig. 6a). Notably, the SAM- and target adenosine-binding sites of METTL3-MTTL14 complex, formed by Gate loop 1 (METTL3 residues 396-410) and Gate loop 2 (METTL3 residues 507-515), are well aligned with the corresponding regions (residues 210-232 and 328-336, respectively) of AMT1 (Supplementary Fig. 6b). Furthermore, a number of DNA-binding sites of AMT1, such as residues Y227, K280 and R357, are conserved in METTL3 (Supplementary Fig. 6c), suggesting that AMT1 and METTL3 may employ a similar protein surface platform for substrate binding. On the other hand, the AMT7 residues R379/R382-correpsonding sites in METTL14 are occluded from solvent access by two α-helices (Supplementary Fig. 6a), suggesting that they are unlikely involved in substrate binding in the METTL3-METTL14 complex. These observations shed light onto the functional conservation and divergence between two evolutionary related methyltransferases.

### Structural modeling analysis of the substrate binding by the AMT1-AMT6 complex

The AMT1-AMT6 complex, in which the AMT7 subunit in AMT1-AMT7 complex is replaced by AMT6, is an alternative methyltransferase in maintaining the 6mA patterns shortly after DNA replication^14^. With reduced methylation activity than the AMT1-AMT7 complex in vitro, the AMT1-AMT6 complex maintains the 6mA DNA methylation in cells in a proceeding cell nuclear antigen (PCNA)-dependent manner^14^. To illustrate the substrate binding by the AMT1-AMT6 complex, we generated a structural model of the AMT1-AMT6 complex based on the structure of AMT1-AMT7-P1-P2 complex and AlphaFold3 prediction^39^. The AMT1-AMT7 and AMT1-AMT6 complexes are structurally similar, with an RMSD of 1.77 Å over 1279 aligned Cα atoms (Supplementary Fig. 7a). Notably, the DNA-binding sites of AMT7, including R379 and R382, are also conserved in AMT6 (Supplementary Fig. 7b), suggesting a conserved DNA-binding mechanism between the two homologues. Nevertheless, AMT6 differs from AMT7 in the sequence of the disordered N-terminus (Supplementary Fig. 7b). In addition, the surface electrostatic potential of AMT6 appears less positively charged than that of AMT7, due to amino acid variation in the region near to the DNA-binding site (Supplementary Fig. 7a-d). Whether such a difference in the DNA-binding site contributes to the differential methylation activity between the AMT1-AMT7 and AMT1-AMT6 complexes awaits further investigation.

## Discussion

6mA DNA methylation in eukaryotes has gained increasing attention due to its functional implication in epigenetic regulation and genomic assembly^40^. This study, through structure determination of an inhibitor-trapped state of the AMT complex from ciliated *Tetrahymena thermophila*, combined with enzymatic analysis, provides a framework for understanding how the AMT complex engages DNA substrate for 6mA maintenance methylation, with important implications in functional regulation of 6mA DNA methylation in eukaryotes.

First of all, this study lends strong evidence that the AMT complex undergoes an open-to-closed conformational transition upon the substrate binding (Fig. 5). Previous structural characterization of the AMT complex reveals that, in the absence of DNA, the AMT1, AMT7, AMTP2 and the C-terminal helix of AMTP1 adopts an open conformation, while the ND domain of AMTP1 is disordered^21,22^. While these studies provide a framework for the functional assembly of the AMT complex, the lack of structural details for the ND domain of AMTP1 raises a question on its functional role in AMT-mediated 6mA DNA methylation. To overcome the challenge of the dynamic AMT-DNA interaction, we took advantage of a DNA-competitive inhibitor, the OCR protein, to stabilize the DNA-binding components of the AMT complex. As a result, we were able to capture a saddle-shaped architecture formed by AMT1, AMT7, AMTP1, and AMTP2, in which AMTP1 and the AMT1-AMT7 heterodimer coordinately flank the OCR protein. Consistently, our in vitro enzymatic analysis reveals that mutations of the putative DNA-binding sites of AMT1, AMT7 and AMTP1 impact the AMT activity to various extents. Our comparative structural analysis further demonstrated that the ND domain of AMTP1 shares a similar functional site with other members of the homeodomain family, implying a role for this domain to target the major groove of DNA substrate. Whether AMTP1 contributes to the DNA sequence specificity of the AMT complex, as what was observed for the TRD domain of a C-5 DNA methyltransferase^18,41^, has yet to be elucidated.

**Fig. 5:**
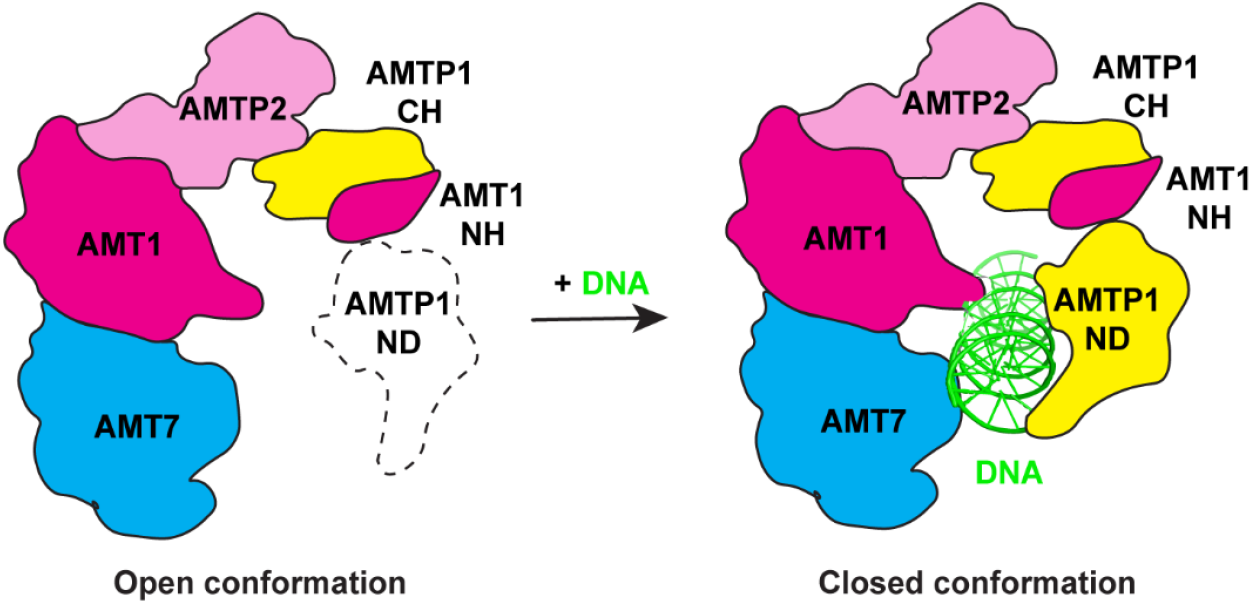
A working model of the AMT complex. In the absence of DNA substrate, the AMTP1 N-terminal domain is dynamically disordered, resulting in an open conformation. In the presence of DNA substrate, the AMTP1 is dynamically stabilized and cooperates with the AMT1-AMT7 dimer in DNA binding, resulting in a closed conformation for processive 6mA methylation.

It has been well known that DNA-templated enzymes often adopt a ring or saddle-shaped structure to process DNA^42^. For instance, encircling of the DNA by eukaryotic DNA polymerase processivity factor PCNA^43^ and bacteriophageT4 gp45^44^ allows the polymerases to move rapidly without dissociation from the template. Similarly, the restriction-modification enzyme M.TaqI from *Thermus aquaticus*, adopts a saddle shape for 6mA DNA methylation^36^, as well as the DNA restriction enzyme *Bam*HI^45^, and the TATA-box binding protein (TBP)^46^ all adopt a saddle conformation to slide along DNA for processive enzymatic or scanning actions. In this context, the substrate binding-induced conformational closure of the AMT complex may underpin its processive methylation along DNA^9,12^. A detailed understanding of the enzymatic kinetics of the AMT complex awaits further investigation.

### Limitation of this study

Through introducing the DNA competitive inhibitor, OCR protein, we were able to capture the closed conformation of the AMT complex that is important for its substrate binding and methylation. This study therefore provides important insights into the AMT complex-mediated processive 6mA DNA methylation. However, the lack of defined density for OCR makes us unable to generate an accurate model for the AMT-OCR interaction. Furthermore, how the DNA substrate undergoes a transition from the encounter state to a productive methylation state remains uncharacterized. Future study on the productive state of the AMT-DNA complex is warranted for a detailed understanding of 6mA DNA methylation by the AMT complex.

## Materials and methods

### Plasmid

The genes for full-length *Tetrahymena thermophila* AMT1 and AMT7 were inserted in tandem into an in-house bacterial expression vector, in which the AMT1 was preceded by an N-terminal His_6_-MBP tag and a TEV cleavage site. Both DNA fragments encoding full-length *Tetrahymena thermophila* AMTP1 and AMTP2 were cloned into a modified pRSF vector preceded by an N-terminal His_6_-SUMO tag and ULP1 (ubiquitin-like protease 1) cleavage site. The DNA fragment encoding *Escherichia* phage T7 OCR (residues 1-117; uniport number. P03775) was chemically synthesized by Integrated DNA Technologies and cloned into the pRSF vector.

### Protein expression and purification

The expression plasmids for AMT1, AMT7, and AMTP2 were co-transformed into BL21 (DE3) RIL cells strain (Agilent Technologies). The cells were initially grown at 37 °C and were shifted to 16°C when OD_600_ reached 1.0. The cells were induced by the addition of 0.13 mM isopropyl β-D-1-thiogalactopyranoside (IPTG) and continued to grow overnight. The cells were lysed in buffer containing 50 mM Tris-HCl (pH 8.0), 1 M NaCl, 25 mM imidazole, 10% glycerol, 10 μg/mL DNase I, and 1 mM PMSF. The fusion proteins were purified using a Ni-NTA affinity column. The AMT1-AMT7-AMTP2 complex was then treated with ULP1 protease to remove the His_6_-SUMO tag and TEV protease to remove the His_6_-MBP tag, followed by ion-exchange chromatography on a HiTrap Heparin HP column (Cytiva) and size-exclusion chromatography on a HiLoad 16/600 Superdex 200 pg column (GE Healthcare) in 20 mM Tris-HCl (pH 7.5), 100 mM NaCl, 5% glycerol, and 5 mM DTT. Purified protein samples were stored at -80 °C for further use.

The AMTP1 and OCR proteins were purified in a similar manner to the above, except that the cells were shifted to 16 °C when OD_600_ reached 0.8. After cell lysis, the AMTP1 or OCR protein was purified using a Ni-NTA affinity column, followed by incubation with ULP1 protease. To remove the His_6_-SUMO tag, the AMTP1 protein was further purified by ion-exchange chromatography on a HiTrap Heparin HP column (Cytiva), and the OCR protein was purified using a HiTrap QHP Sepharose column (Cytiva). The tag-free AMTP1 or OCR protein was finallypurified by size exclusion chromatography on a HiLoad 16/600 Superdex 75 pg column (GE Healthcare) in 20 mM Tris-HCl (pH 7.5), 100 mM NaCl, 5% glycerol and 5 mM DTT. Purified protein samples were stored at -80 °C before use. The mutants of AMT1, AMT7 and AMTP1 were generated by site-directed mutagenesis and purified in the same manner as wild-type (WT) proteins.

### Radioactivity-based DNA methylation assay

To assess the OCR-mediated inhibition of the AMT complex, a synthesized 12-mer DNA duplex containing a single hemimethylated ApT site (Upper strand: 5’-GCAAGXTCAACG-3’, X=6-methyladenine; Lower strand: 5’-CGTTGATCTTGC -3’) was used as DNA substrate. The 20-µL reaction mixture contained 1 µM AMT1-AMT7-AMTP1-AMTP2 complex, 1 µM DNA and 0.56 µM S-adenosyl-L-[methyl-^3^H] methionine with a specific activity of 18 Ci/mmol (PerkinElmer) in buffer containing 50 mM Tris-HCl (pH 8.0), 100 mM NaCl, 200 µg/mL BSA, 0.05% β-mercaptoethanol, and 5% glycerol in the presence or absence of 2 µM OCR. The reaction was performed at 30 °C for 30 minutes before being quenched by 5 µL of 10 mM cold SAM. Eight μL of the mixtures were loaded onto Hybond N nylon membrane (GE Healthcare) and dried out at room temperature, followed by washing with 0.2 M ammonium bicarbonate (pH 8.2) twice, deionized water once, and 95% ethanol once. The dry membrane was then transferred into a vial containing 3-mL scintillation buffer (Fisher) and read by a Beckman LS6500 counter. The assays were performed in triplicate.

### Luminescence-based DNA methylation assay

For mutational analysis of the AMT complex, the MTase-Glo^TM^ Methyltransferase Assay kit was used to detect activity (Promega). Specifically, 1 μM WT or mutant AMT complex was mixed with 10 μM SAM and the aforementioned hemimethylated 12-mer DNA duplex or unmethylated, self-complimentary 24-mer DNA duplex (5’-CGCGATCGATGATCATCGATCGCG-3’) in a buffer containing 20 mM Tris-HCl (pH 8.0), 50 mM NaCl, 1 mM EDTA, 3 mM MgCl_2_, 100 µg/mL BSA, and 1 mM DTT at 30 °C. The reaction lasted for 30 minutes before being quenched by 0.1% trifluoroacetic acid. Subsequently, the reaction mixture was first mixed with 2 μL 6× MTase-Glo^TM^ reagent at room temperature for 30 minutes and then with 12 μL MTase-Glo^TM^ detection solution for 30 minutes. The luminescence of the samples was measured in an opaque white 384-well microwell plate (Revvity) by CLARIOstar Plus (BMG LABTECH) at 25 °C.

### Analytical size-exclusion chromatography

A 400-μL sample containing 18 µM AMT1-AMT7-AMTP1-AMTP2 complex mixed with 54 µM OCR was loaded onto Superdex 200 increase 10/300 GL column (GE Healthcare) pre-equilibrated with 20 mM Tris-HCl (pH 7.5), 100 mM NaCl, 5% glycerol, and 5 mM DTT. The proteins in each of the elution fractions were visualized by the SDS-PAGE imaging.

### Cryo-EM sample preparation and data collection

The concentration of the AMT1-AMT7-AMTP1-AMTP2-OCR complex was adjusted to 1.1 mg/mL in a buffer containing 20 mM Tris-HCl (pH 7.5), 100 mM NaCl, 5 mM DTT, and 2.5% Glycerol. Three μl of the complex sample was applied to Quantifoil® (1.2/1.3) grids, which were glow discharged at 15 mA for 90 s. The grids were blotted for 7 s at 95% humidity with blot force 2 and plunge frozen using a Vitrobot Mark IV (Thermo Fisher).

The cryo-EM grids were imaged on a 300 keV Titan Krios cryo-electron microscope equipped with a K3 camera (Gatan) by the Nation Center for CryoEM Access and Training (NCCAT). Micrographs were collected at a magnification of ×81,000 with a super-resolution pixel size of 0.53 Å. The exposures were set with a total dose of ∼54 e^-^/Å^2^ over 50 frames and a defocus range of -0.4 μm to -3.1 μm.

### Cryo-EM data processing

In total 5137 movies were processed using the CryoSPARC software package (v4.01)^47^. All movies are subject to 2×2 binning, patch motion correction and patch-based contrast transfer function estimation. Subsequently, 2.6 million particles were picked using the Topaz program^48^, and extracted with a downscaled pixel size of 2.12 Å per pixel. After several rounds of 2D classification, 0.2 million particles were selected for 3D ab initio reconstruction and subsequent heterogeneous refinement. The particles with visible features of secondary and tertiary structures were re-extracted with pixel size of 1.06 Å per pixel. The final set of particles were subject to removal of duplicates and non-uniform refinement, resulting in a final resolution of 3.59 Å, as given by the Fourier shell correlation criterion (FSC 0.143).

### Model building and refinement

The structural model for AMT1-AMT7-AMTP2 complex was derived from that of the reported apo-state AMT complex (PDB 7YI8)^22^. The structural models for AMTP1 and OCR were generated by AlphaFold3 method^39^. Fitting the models into the density map was carried out using UCSF Chimera (v1.16)^49^ and ChimeraX (v1.6.1)^50^. The final structural model was obtained after iterative model building in Coot (v0.8.9.1)^51^, followed by real-space refinement in Phenix^52^.

## Acknowledgments

This work was supported by NIH grant (R35GM119721 to J.S.) and NSF award (MCB-2435178 to Y.L.). Cryo-EM data were collected at the National Center for CryoEM Access and Training (NCCAT) and the Simons Electron Microscopy Center located at the New York Structural Biology Center, supported by the NIH Common Fund Transformative High Resolution Cryo-Electron Microscopy program (U24 GM129539,) and by grants from the Simons Foundation (SF349247) and NY State Assembly.

## Author Contributions

Z.S., S.Y., J.L., and P.A. performed the experiments, Y.L. and J.S. conceived and oversaw the study. Z.S. and J.S. wrote the manuscript and all authors approved the manuscript.

## Competing Interest

The authors declare no competing interest.

## Data availability

All data needed to evaluate the conclusions in the paper are present in the paper and/or the Supplementary Materials. Coordinates for the AMT1-AMT7-AMTP1-AMTP2-OCR complex have been deposited in the Protein Data Bank under accession codes 9O6K. The cryo-EM density map has been deposited to the EMDB under the accession numbers of EMD-70174.

## Notes

### Competing Interest Statement

The authors have declared no competing interest.

