## Supplementary information for "Structural insight into the substrate binding of the AMT complex via an inhibitor-trapped state"

[2]Department of Biochemistry and Molecular Medicine, University of Southern California, Los Angeles, CA 90033, USA

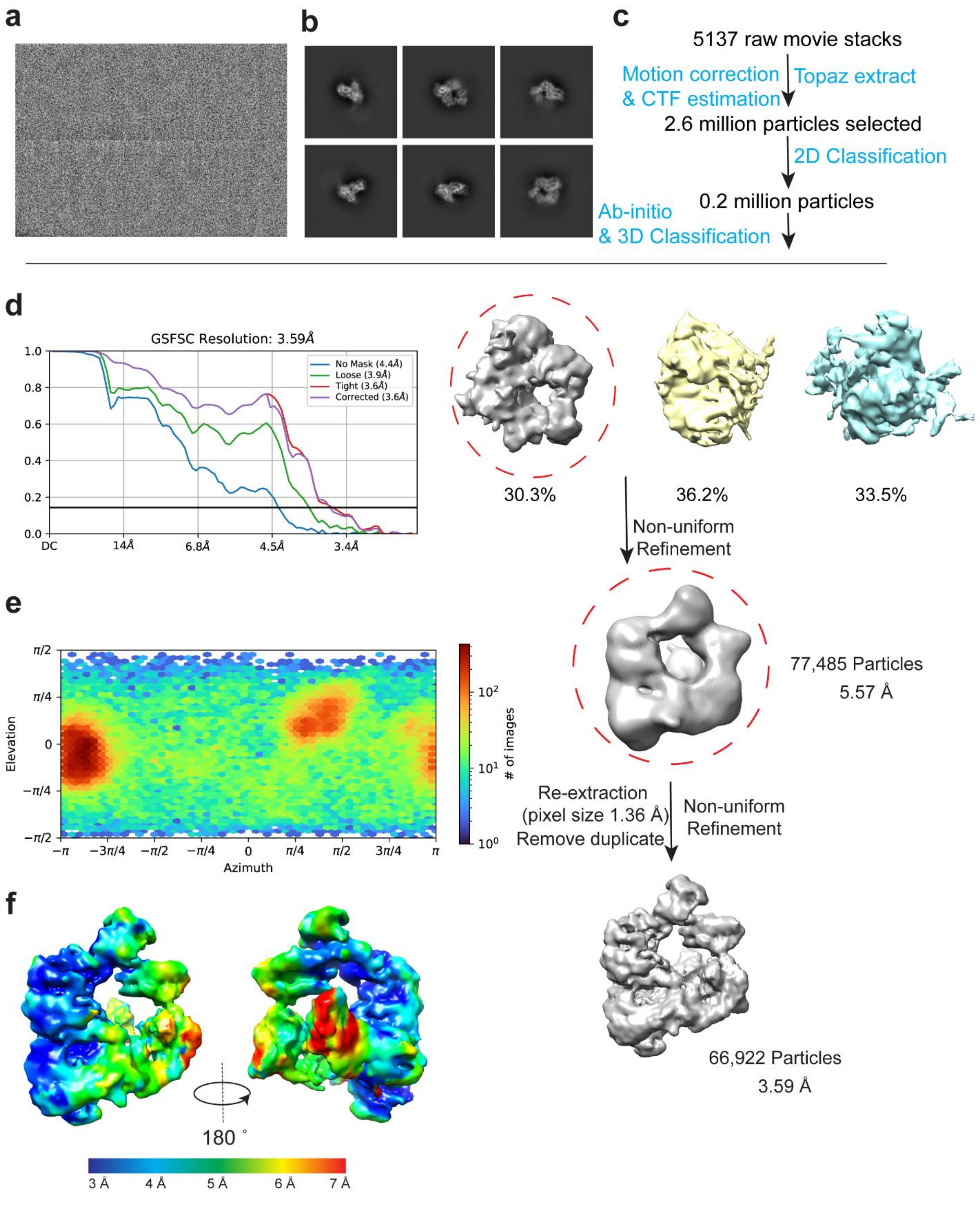


**Supplementary Fig. 1. Data processing workflow for cryo-EM reconstruction of the AMT-OCR complex. a** A representative micrograph of the AMT-OCR complex. **b** Representative 2D classes of the AMT-OCR complex from CryoSPARC. **c** Flow chart of the cryo-EM data processing. **d** Fourier shell correlation (FSC) curve of the AMT-OCR complex map as a function of resolution using CryoSPARC output. **e** Angular distribution calculated in CryoSPARC for particle projections. Heat map shows number of particles for each viewing angle. **f** Local resolution map of the AMT-OCR complex.

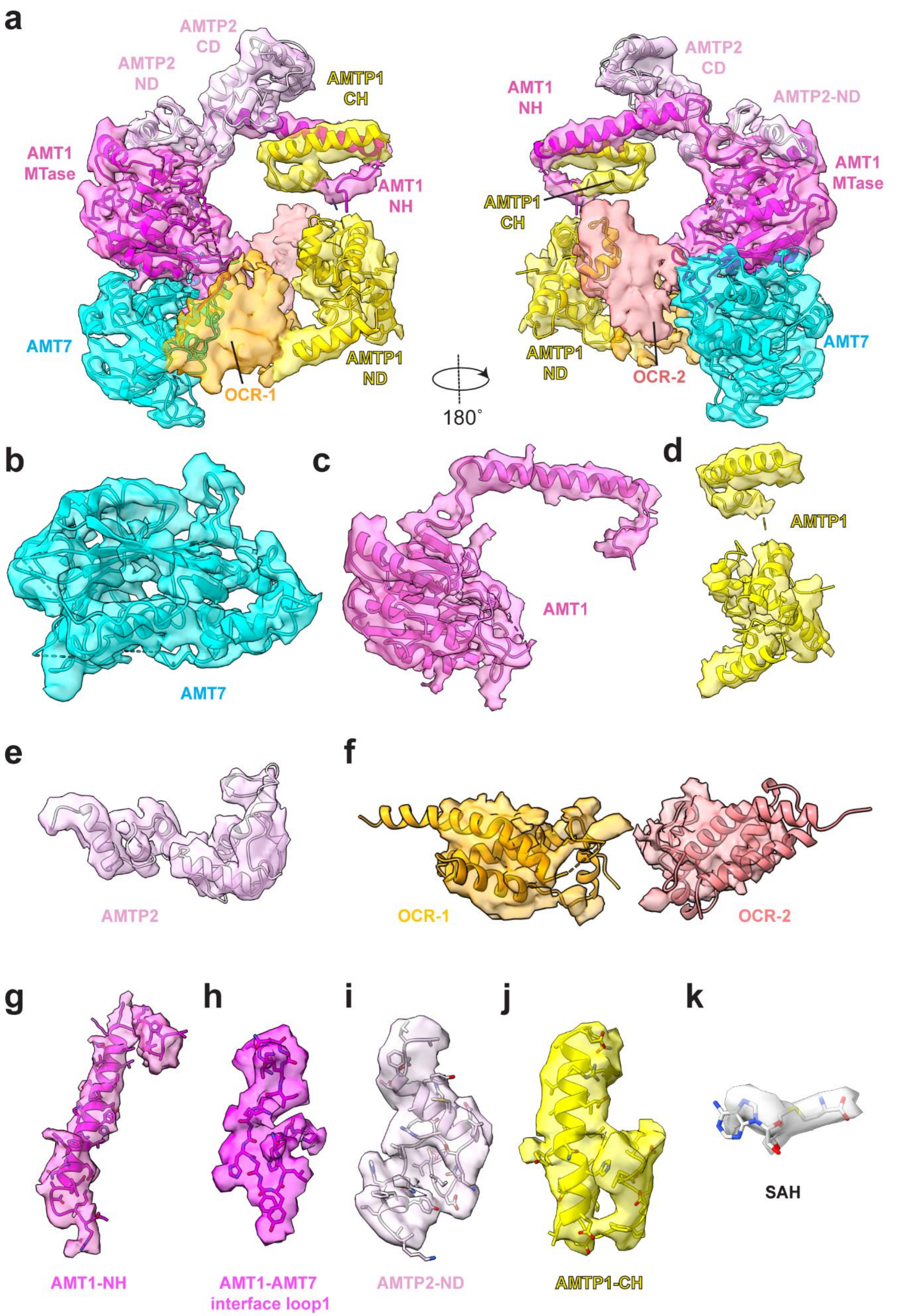
a
AMTP2
CD
AMTP2
ND
AMTP1
CH
AMT1
MTase
AMT1
NH
AMT7
OCR-1
AMTP1
ND
180˚
AMT1
NH
AMTP2
CD
AMTP2-ND
AMT1
MTase
AMTP1
CH
AMTP1
ND
OCR-2
AMT7
b
AMT7
c
AMT1
d
AMTP1
e
AMTP2
f
OCR-1
OCR-2
g
AMT1-NH
h
AMT1-AMT7
interface loop1
i
AMTP2-ND
j
AMTP1-CH
k
SAH

**Supplementary Fig. 2. Cryo-EM density maps for the AMT-OCR complex. a-k** Density map and atomic model of the AMT-OCR complex (a), AMT7 (b), AMT1 (c), AMTP1 (d), AMTP2 (e), OCR dimer (f), the AMT1 N-terminal helix (AMT1-NH) (g), the AMT1-AMT7 interface loop (residues 282-299) (h), the AMTP2 N-terminal domain (AMTP2-ND) (i), the AMTP1 C-terminal helix (AMTP1-CH) (j), and SAH molecule (k). The model for OCR dimer was not included in the final model due to relatively weak density.

**a**

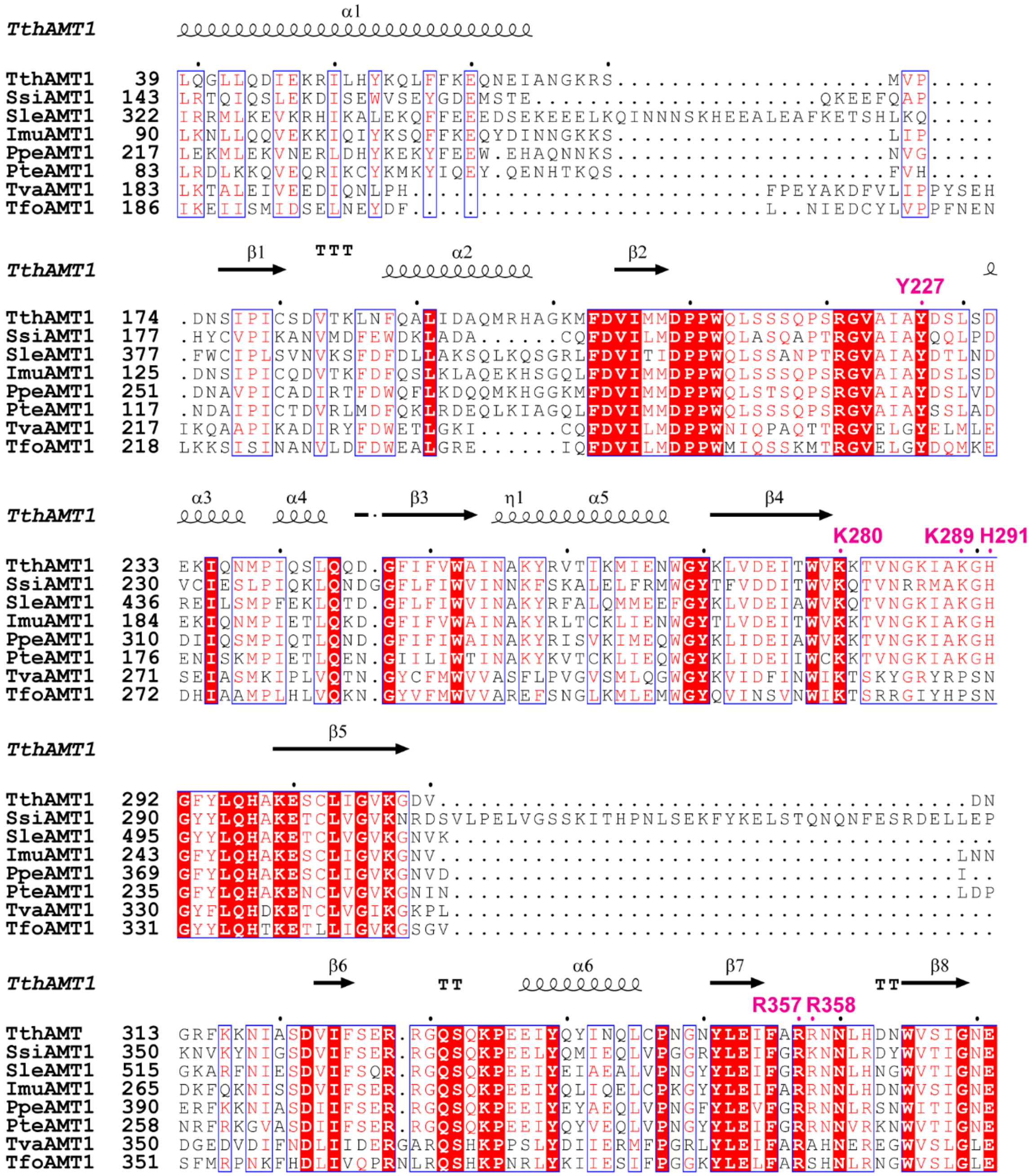
TthAMT1
α1
TthAMT1 39 LQGLLQDIEKRILHYKQLFFKEQNEIANGKRS...................MVP.....
SsiAMT1 143 LRTQIQSLEKDISEWVSEYGDEMSTE..................QKEEFQAP.....
SleAMT1 322 IRRMLKEVKRHIKALEKQFFEEEDSEKEEELKQINNNSKHEEALEAFKETSHLKQ.....
ImuAMT1 90 LKNLLQQVEKKIQIYKSQFFKEQYDINNGKKS...................LIP.....
PpeAMT1 217 LEKMLEKVNERLDHYKEKYFEEW.EHAQNNKS...................NVG.....
PteAMT1 83 LRDLKKQVEQRIKCYKMKYIQEY.QENHTKQS...................FVH.....
TvaAMT1 183 LKTALEIVEEDIQNLPH.........................FPEYAKDFVLIPPYSEH
TfoAMT1 186 IKEIISMIDSELNEYDF..........................L..NIEDCYLVPPFNEN
TthAMT1
β1 TTT α2 β2
Y227
TthAMT1 174 .DNSIPICSDVTKLNFQALIDAQMRHAGKMFDVIMMDPPWQLSSSQPSRGVAIAYDSLSD
SsiAMT1 177 .HYCVPIKANVMDFEWDKLADA......CQFDVILMDPPWQLASQAPTRGVAIAYQQLPD
SleAMT1 377 .FWCIPLSVNVKSFDFDLLAKSQLKQSGRLFDVITIDPPWQLSSANPTRGVAIAYDTLND
ImuAMT1 125 .DNSIPICQDVTKFDFQSLKLAQEKHSGQLFDVIMMDPPWQLSSSQPSRGVAIAYDSLSD
PpeAMT1 251 .DNAVPICADIRTFDWQFLKDQQMKHGGKMFDVIMMDPPWQLSTSQPSRGVAIAYDSLVD
PteAMT1 117 .NDAIPICTDVRLMDFQKLRDEQLKIAGQLFDVIMMDPPWQLSSSQPSRGVAIAYSSLAD
TvaAMT1 217 IKQAAPIKADIRYFDWETLGKI......CQFDVILMDPPWNIQPAQTTRGVELGYELMLE
TfoAMT1 218 LKKSISINANVLDFDWEALGRE......IQFDVILMDPPWMIQSSKMTRGVELGYDQMKE
TthAMT1
α3 α4 β3 η1 α5 β4
K280 K289 H291
TthAMT1 233 EKIQNMPIQSLQQD.GFIFVWAINAKYRVTIKMIENWGYKLVDEITWVKKTVNGKIAKGH
SsiAMT1 230 VCIESLPIQKLQNDGGFLFIWVINNKFSKALELFRMWGYTFVDDITWVKQTVNRRMAKGH
SleAMT1 436 REILSMPFEKLQTD.GFLFIWVINAKYRFALQMMEEFGYKLVDEIAWVKQTVNGKIAKGH
ImuAMT1 184 EKIQNMPIETLQKD.GFIFVWAINAKYRLTCKLIENWGYTLVDEITWVKKTVNGKIAKGH
PpeAMT1 310 DIIQSMPIQTLQND.GFIFIWAINAKYRISVKIMEQWGYKLIDEIAWVKKTVNGKIAKGH
PteAMT1 176 ENISKMPIETLQEN.GIILIWTINAKYKVTCKLIEQWGYKLIDEIIWCKKTVNGKIAKGH
TvaAMT1 271 SEIASMKIPLVQTN.GYCFMWVVASFLPVGVSMLQGWGYKVIDFINWIKTSKYGRYRPSN
TfoAMT1 272 DHIAAMPLHLVQKN.GYVFMWVVAREFSNGLKMLEMWGYQVINSVNWIKTSRRGIYHPSN
TthAMT1
β5
TthAMT1 292 GFYLQHAKESCLIGVKGDV.......................................DN
SsiAMT1 290 GYYLQHAKETCLVGVKNRDSVLPELVGSSKITHPNLSEKFYKELSTQNQNFESRDELLEP
SleAMT1 495 GYYLQHAKETCLVGVKGNVK........................................
ImuAMT1 243 GFYLQHAKESCLIGVKGNV.......................................LNN
PpeAMT1 369 GFYLQHAKESCLIGVKGNVD......................................I..
PteAMT1 235 GFYLQHAKENCLVGVKGNIN......................................LDP
TvaAMT1 330 GYFLQHDKETCLVGIKGKPL........................................
TfoAMT1 331 GYYLQHTKETLLIGVKGSGV........................................
TthAMT1
β6 TT α6 β7 TT β8
R357 R358
TthAMT 313 GRFKKNIASDVIFSER.RGQSQKPEEIYQYINQLCPNGNYLEIFARRNNLHDNWVSIGNE
SsiAMT1 350 KNVKYNIGSDVIFSER.RGQSQKPEELYQMIEQLVPGGRYLEIFGRKNNLRDYWVTIGNE
SleAMT1 515 GKARFNIESDVIFSQR.RGQSQKPEEIYEIAEALVPNGYYLEIFGRRNNLHNGWVTIGNE
ImuAMT1 265 DKFQKNISSDVIFSER.RGQSQKPEEIYQLIQELCPKGYYLEIFARRNNLHDNWVSIGNE
PpeAMT1 390 ERFKKNIASDIIFSER.RGQSQKPEEIYEYAEQLVPNGFYLEVFGRRNNLRSNWITIGNE
PteAMT1 258 NRFRKGVASDIIFSER.RGQSQKPEEIYQYVEQLVPNGYYLEIFGRRNNVRKNWVTIGNE
TvaAMT1 350 DGEDVDIFNDLIIDERGARQSHKPPSLYDIIERMFPGRLYLEIFARAHNEREGWVSLGLE
TfoAMT1 351 SFMRPNKFHDLIVQPRNLRQSHKPNRLYKIIESIFPGGKYLELFARSHNLRNGWVSLGLE

**b**

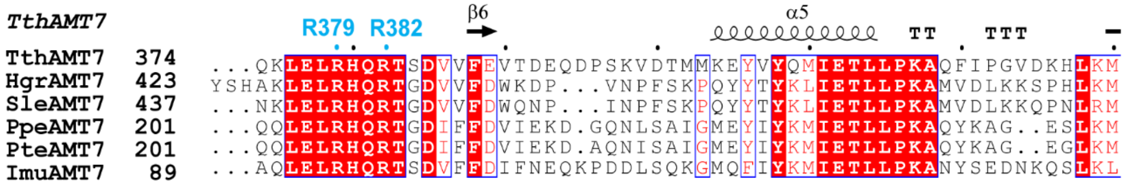
TthAMT7
R379 R382
β6 α5 TT TTT
TthAMT7 374 ...QKLELRHQRTSDVVFEVTDEQDPSKVDTMMKEYVYQMIETLLPKAQFIPGVDKHLKM
HgrAMT7 423 YSHAKLELRHQRTGDVVFDWKDP...VNPFSKPQYYTYKLIETLLPKAMVDLKKSPHLKM
SleAMT7 437 ...NKLELRHQRTGDVVFDWQNP...INPFSKPQYYTYKLIETLLPKAMVDLKKQPNLRM
PpeAMT7 201 ...QQLELRHQRTGDIFFDVIEKD.GQNLSAIGMEYIYKMIETLLPKAQYKAG..ESLKM
PteAMT7 201 ...QQLELRHQRTGDIFFDVIEKD.AQNISAIGMEYIYKMIETLLPKAQYKAG..EGLKM
ImuAMT7 89 ...AQLELRHQRTSDVFFDIFNEQKPDDLSQKGMQFIYKMIETLLPKANYSEDNKQSLKL

**Supplementary Fig. 3. Sequence alignment of AMT1 and AMT7 from various species. a** Sequence alignment of AMT1 proteins. TthAMT1: *Tetrahymena thermophila* AMT1 (XP_001032074), SsiAMT1: *Smittium simulii* AMT1 (PVU88469), SleAMT1: *Stylonychia lemnae* AMT1 (CDW73011), ImuAMT1: *Ichthyophthirius multifiliis* AMT1 (XP_004035673), PpeAMT1: *Pseudocohnilembus persalinus* AMT1 (KRX04523), PteAMT1: *Paramecium tetraurelia* AMT1 (XP_001457964), TvaAMT1: *Trichomonas vaginalis G3* AMT1 (EAX86932), TfoAMT1: *Tritrichomonas foetus* AMT1 (OHT14409). Identical or similar residues are boxed and colored red. The secondary structures corresponding to TthAMT1 are indicated on top. The OCR-contact sites of AMT1 are indicated in magenta. **b** Sequence alignment of AMT1 proteins. TthAMT7: *Tetrahymena thermophila* AMT7 (7YI8_A), HgrAMT7: *Halteria grandinella* AMT7 (TNV81034), SleAMT7: *Stylonychia lemnae* AMT7 (CDW82373), PpeAMT7: *Paramecium pentaurelia* AMT7 (CAD8164721), PteAMT7: *Paramecium tetraurelia* AMT7 (XP_001442122), ImuAMT7: *Ichthyophthirius multifiliis* AMT7 (XP_004030651), TthAMT6: *Tetrahymena thermophila* AMT6 (XP_001010335). Identical or similar residues are boxed and colored red. Completely conserved residues are shaded in red. The secondary structures corresponding to TthAMT7 are indicated on top. The OCR-contact sites of AMT7 are indicated in cyan.

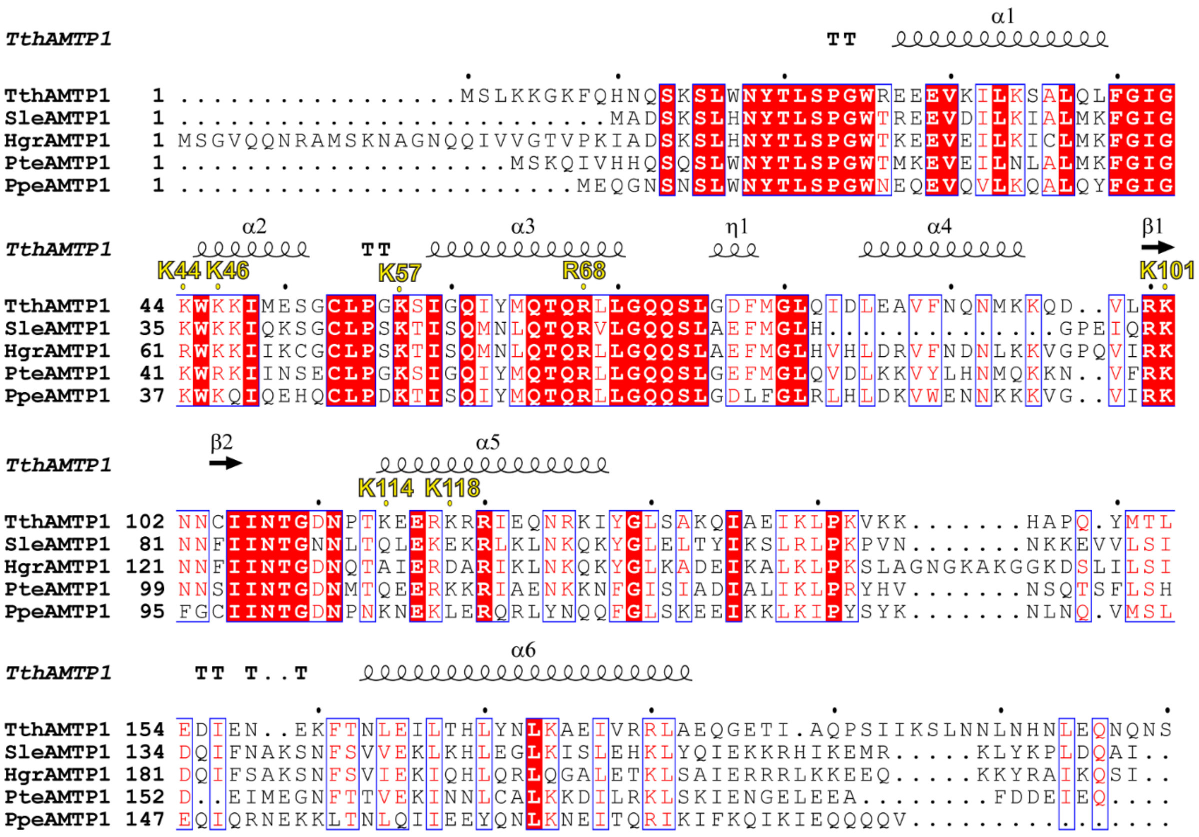


**Supplementary Fig. 4. Sequence alignment of AMTP1 from various species.** TthAMTP1: *Tetrahymena thermophila* AMTP1 (XP_001009903), SleAMTP1: *Stylonychia lemnae* AMTP1 (CDW84997), HgrAMTP1: *Halteria grandinella* AMTP1 (TNV87513), PteAMTP1: *Paramecium tetraurelia* AMTP1 (XP_001443306), PpeAMTP1: *Pseudocohnilembus persalinus* AMTP1 (KRX10274). The OCR-contact residues of AMTP1 are labeled in yellow.

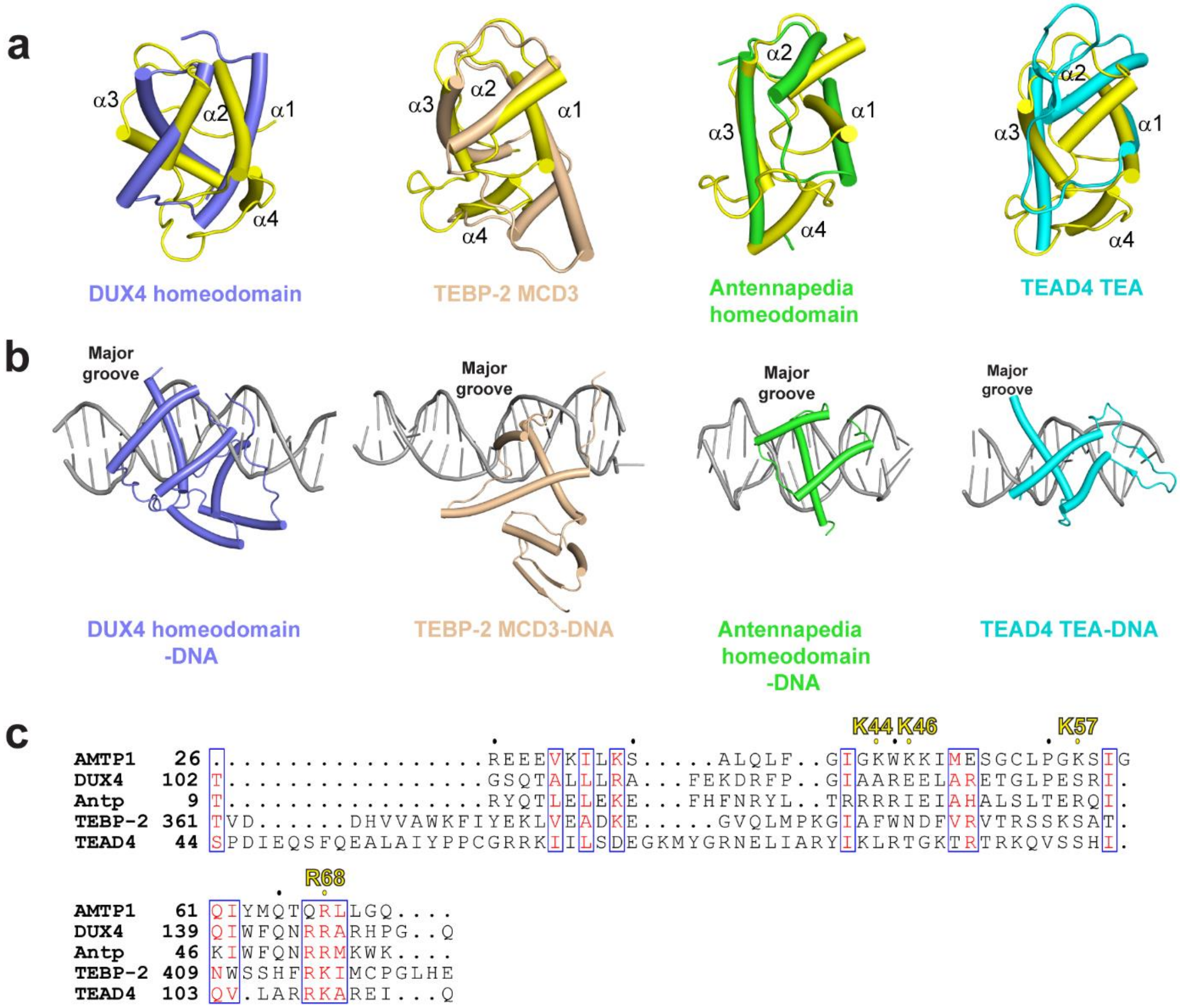


**Supplementary Fig. 5. Structural comparison of AMTP1 and other homodomain proteins.** **a** Structural alignment of AMTP1 homeodomain (yellow) with DNA-bound homeodomain-containing proteins: DUX4 homeodomain (PDB 6E8C, slate), TEBP-2 MCD3 (PDB 8U8L, wheat), Antennapedia homeodomain (PDB 9ANT, green), and TEAD4 TEA (PDB 5GZB, cyan). The long C-terminus of α3 in TEAD4 TEA was removed for clarity. **b** Structure of the homeodomain-containing proteins in complex with DNA: DUX4 homeodomain (slate)-DNA (grey) (PDB 6E8C), TEBP-2 MCD3 (wheat)-DNA (grey) (PDB 8U8L), Antennapedia homeodomain (green)-DNA (grey) (PDB 9ANT), and TEAD4 TEA (cyan)-DNA (grey) (PDB 5GZB). **c** Sequence alignment of AMPT1 homeodomain with that of DUX4, Antennapedia (Antp), TEBP-2, and TEAD4. The OCR-contact sites of AMTP1 are indicated on top.

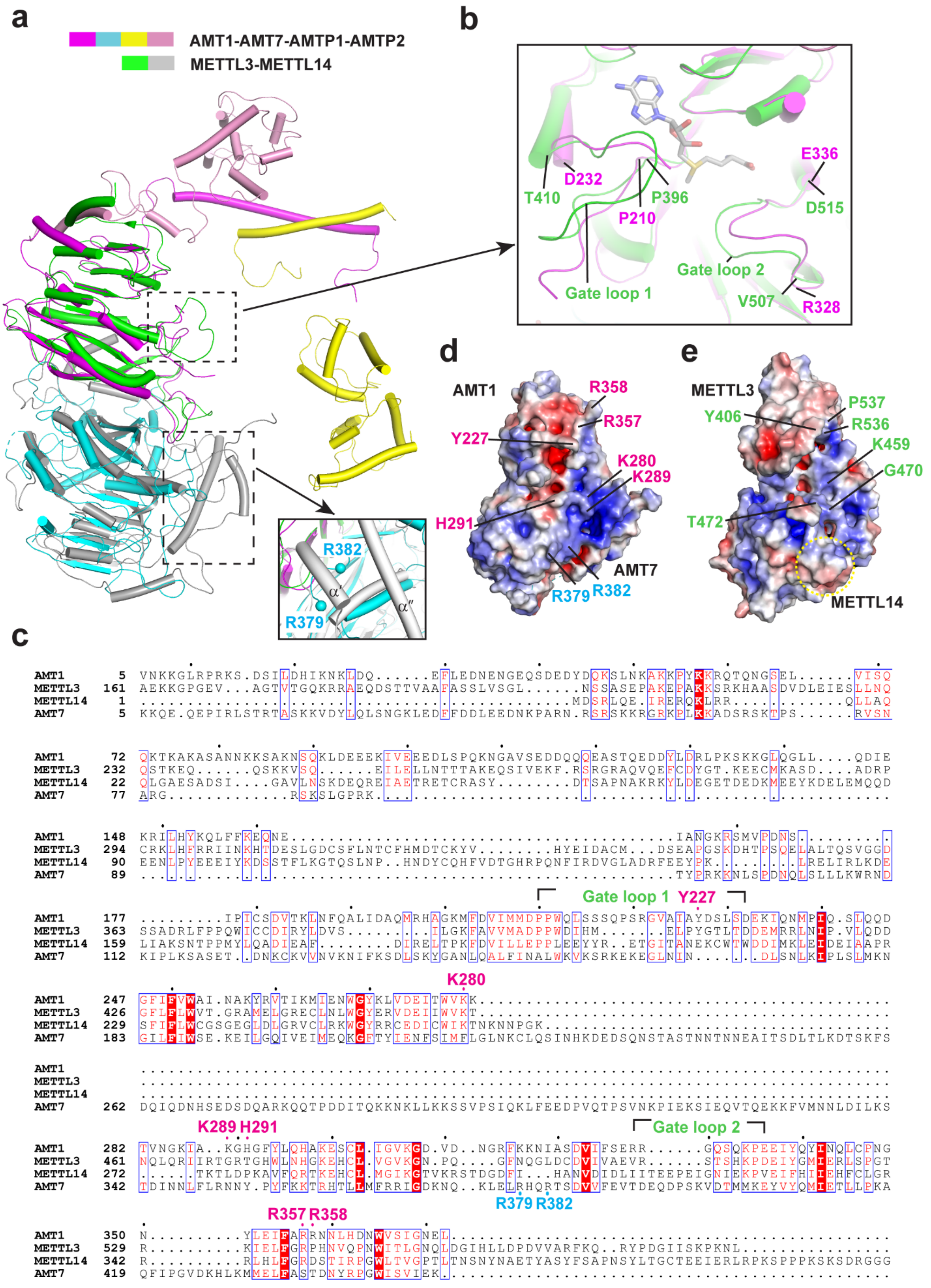

a
AMT1-AMT7-AMTP1-AMTP2
METTL3-METTL14
R382
R379
α′
α″
b
T410
D232
P396
P210
E336
D515
Gate loop 1
Gate loop 2
V507
R328
d
AMT1
R358
R357
Y227
K280
K289
H291
AMT7
R379
R382
e
METTL3
P537
Y406
R536
K459
G470
T472
METTL14
c
AMT1 5 VNKKGLRPRKS.DSILDHIKNKLDQ......EFLEDNENGEQSDEDYDQKSLNKAKKPYKKRQTQNGSEL......VISQ
METTL3 161 AEKKGPGEV...AGTVTGQKRRAEQDSTTVAAFASSLVSGL......NSSASEPAKEPAKKSRKHAASDVDLEIESLLNQ
METTL14 1 ...............................................MDSRLQE.IRERQKLRR...........QLLAQ
AMT7 5 KKQE.QEPIRLSTRTASKKVDYLQLSNGKLEDFFDDLEEDNKPARN.RSRSKKRGRKPLKKADSRSKTPS......RVSN
AMT1 72 QKTKAKASANNKKSAKNSQKLDEEEKIVEEEDLSPQKNGAVSEDDQQQEASTQEDDYLDRLPKSKKGLQGLL....QDIE
METTL3 232 QSTKEQ......QSKKVSQ......EILELLNTTTAKEQSIVEKF.RSRGRAQVQEFCDYGT.KEECMKASD....ADRP
METTL14 22 QLGAESADSI....GAVLNSKDEQREIAETRETCRASY........DTSAPNAKRKYLDEGETDEDKMEEYKDELEMQQD
AMT7 77 ARG..............RSKSLGPRK.........................................................
AMT1 148 KRILHYKQLFFKEQNE..................................IANGKRSMVPDNS..........
METTL3 294 CRKLHFRRIINKHTDESLGDCSFLNTCFHMDTCKYV........HYEIDACM...DSEAPGSKDHTPSQELALTQSVGGD
METTL14 90 EENLPYEEEIYKDSSTFLKGTQSLNP..HNDYCQHFVDTGHRPQNFIRDVGLADRFEEYPK.........LRELIRLKDE
AMT7 89 ..............................................TYPRKKNLSPDNQLSLLLKWRND
Gate loop 1
Y227
AMT1 177 .........IPICSDVTKLNFQALIDAQMRHAGKMFDVIMMDPPWQLSSSQPSRGVAIAYDSLSDEKIQNMPIQ.SLQQD
METTL3 363 SSADRLFPPQWICCDIRYLDVS........ILGKFAVVMADPPWDIHM.......ELPYGTLTDDEMRRLNIP.VLQDD
METTL14 159 LIAKSNTPPMYLQADIEAF........DIRELTPKFDVILLEPPLEEYYR..ETGITANEKCWTWDDIMKLEIDEIAAPR
AMT7 112 KIPLKSASET..DNKCKVVNVKNIFKSDLSKYGANLQALFINALWKVKSRKEKEGLNIN.......DLSNLKIPLSLMKN
K280
AMT1 247 GFIFVWAI.NAKYRVTIKMIENWGYKLVDEITWVKK............................................
METTL3 426 GFLFLWVT.GRAMELGRECLNLWGYERVDEIIWVKT............................................
METTL14 229 SFIFLWCGSGEGLDLGRVCLRKWGYRRCEDICWIKTNKNNPGK.....................................
AMT7 183 GILFIWSE.KEILGQIVEIMEQKGFTYIENFSIMFLGLNKCLQSINHKDEDSQNSTASTNNTNNEAITSDLTLKDTSKFS
AMT1 ................................................................................
METTL3 ................................................................................
METTL14 ................................................................................
AMT7 262 DQIQDNHSEDSDQARKQQTPDDITQKKNKLLKKSSVPSIQKLFEEDPVQTPSVNKPIEKSIEQVTQEKKFVMNNLDILKS
K289 H291
Gate loop 2
AMT1 282 TVNGKIA..KGHGFYLQHAKESCL.IGVKGD.VD..NGRFKKNIASDVIFSERR......GQSQKPEEIYQYINQLCPNG
METTL3 461 NQLQRIIRTGRTGHWLNHGKEHCL.VGVKGN.PQ....GFNQGLDCDVIVAEVR......STSHKPDEIYGMIERLSPGT
METTL14 272 ......TKTLDPKAVFQRTKEHCL.MGIKGTVKRSTDGDFI...HANVDIDLIITEEPEIGNIEKPVEIFHIIEHFCLGR
AMT7 342 TDINNLFLRNNY.PYFKKTRHTLLMFRRIGDKNQ..KLELRHQRTSDVVFEVTDEQDPSKVDTMMKEYVYQMIETLLPKA
R379 R382
R357 R358
AMT1 350 N..........YLEIFARRNNLHDNWVSIGNEL...............................................
METTL3 529 R..........KIELFGRPHNVQPNWITLGNQLDGIHLLDPDVVARFKQ..RYPDGIISKPKNL...............
METTL14 342 R..........RLHLFGRDSTIRPGWLTVGPTLTNSNYNAETYASYFSAPNSYLTGCTEEIERLRPKSPPPKSKSDRGGG
AMT7 419 QFIPGVDKHLKMMELFASTDNYRPGWISVIEK................................................

**Supplementary Fig. 6. Structural comparison of OCR-bound AMT complex with METTL3-METTL14 complex. a** Structural alignment of OCR-bound AMT complex with METTL3-METTL14 complex (PDB 5IL1), with the cofactor-binding site boxed and the AMT7 R379- and R382-corresponding region highlighted in the expanded view. Note that the corresponding region in METTL3-METTL14 is shielded by a pair of helices ($\alpha'$ and $\alpha''$). **b** Close-up view of the cofactor-binding Gate loops, with residue numbers delimited. The SAM molecule bound to METTL3-METTL14 complex is shown in stick representation. **c** Sequence alignment of *Tetrahymena* AMT1, human METTL3, human METTL14 and *Tetrahymena* AMT7. Identical or similar residues are boxed and colored red. Completely conserved residues are shaded in red. The potential DNA-binding sites for AMT1 and AMT7 are marked above and below the alignment respectively. **d, e** Electrostatic surfaces of the AMT1-AMT7 (d) and METTL3-METTL14 (e) complexes. The OCR-contact sites of AMT1 and AMT7 are labeled in magenta and cyan, respectively. The corresponding sites in METTL3 are labeled in green. The METTL14 region corresponding to AMT7 R379 and R382, which is occluded from solvent access, are marked by dotted circle.

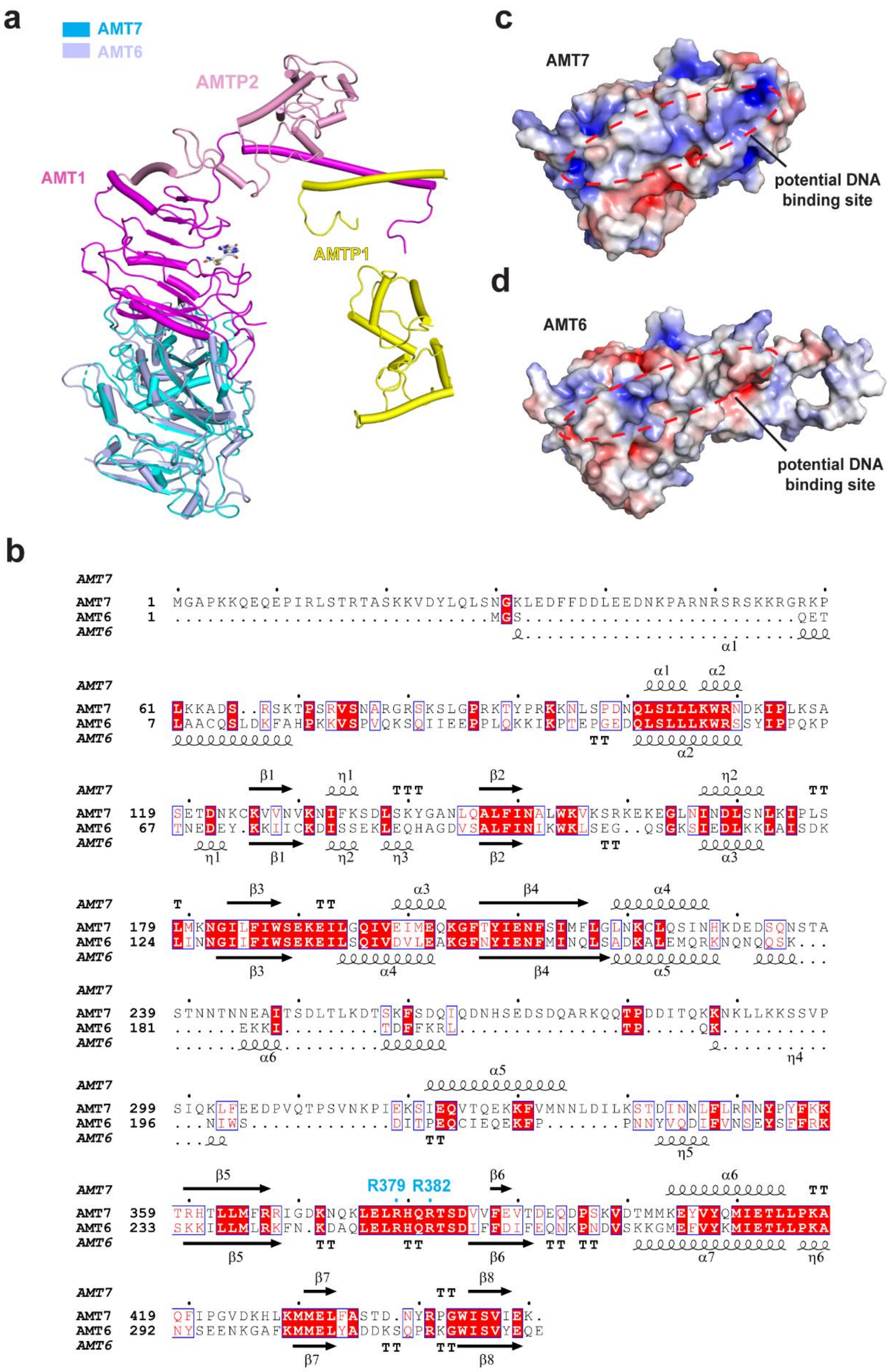

a
AMT7
AMT6
AMTP2
AMT1
AMTP1
c
AMT7
potential DNA binding site
d
AMT6
potential DNA binding site
b
AMT7 1 MGAPKKQEQEPIRLSTRTASKKVDYLQLSNGKLEDFFDDLEEDNKPARNRSRSKKRGRKP
AMT6 1 ............................MGS.....................QET
AMT7 61 LKKADS..RSKTPSRVSNARGRSKSLGPRKTYPRKKNLSPDNQLSLLLKWRNDKIPLKSA
AMT6 7 LAACQSLDKFAHPKKVSPVQKSQIIEEPPLQKKIKPTEPGEDQLSLLLKWRSSYIPPQKP
AMT7 119 SETDNKCKVVNVKNIFKSDLSKYGANLQALFINALWKVKSRKEKEGLNINDLSNLKIPLS
AMT6 67 TNEDEY.KKIICKDISSEKLEQHAGDVSALFINIKWKLSEG..QSGKSIEDLKKLAISDK
AMT7 179 LMKNGILFIWSEKEILGQIVEIMEQKGFTYIENFSIMFLGLNKCLQSINHKDEDSQNSTA
AMT6 124 LINNGIIFIWSEKEILSQIVDVLEAKGFNYIENFMINQLSADKALEMQRKNQNQQSK...
AMT7 239 STNNTNNEAITSDLTLKDTSKFSDQIQDNHSEDSDQARKQQTPDDITQKKNKLLKKSSVP
AMT6 181 ......EKKI.........TDFKRL...............TP.....QK..........
AMT7 299 SIQKLFEEDPVQTPSVNKPIEKSIEQVTQEKKFVMNNLDILKSTDINNLFLRNNYPYFKK
AMT6 196 ...NIWS............DITPEQCIEQEKFP........PNNYVQDIFVNSEYSFFRK
R379 R382
AMT7 359 TRHTLLMFRRIGDKNQKLELRHQRTSDVVFEVTDEQDPSKVDTMMKEYVYQMIETLLPKA
AMT6 233 SKKILLMLRKFN.KDAQLELRHQRTSDIFFDIFEQNKPNDVSKKGMEFVYKMIETLLPKA
AMT7 419 QFIPGVDKHLKMMELFASTD.NYRPGWISVIEK.
AMT6 292 NYSEENKGAFKMMELYADDKSQPRKGWISVYEQE

**Supplementary Fig. 7. Structural and sequence analysis of AMT7 and AMT6**. **a** Structural overlay of AMT7 (cyan) and AMT6 (light blue) in the context of the AMT complex. The structural model for AMT6 was predicted by AlphaFold3. **b** Sequence comparison of AMT6 and AMT7, with the AMT7 residues involved in OCR binding labeled in cyan on top. **c** Electrostatic surface view of AMT7, with the potential DNA-binding surface circled by red dashed line. **d** Electrostatic surface view of AMT6, with the potential DNA-binding site circled by red dashed line.

**Supplementary Table 1. Cryo-EM data collection, refinement and validation statistics.**

| **State** | **AMT1-AMT7-AMTP1-AMTP2** |
|---|---|
| **Code** | (EMD-70174, PDB 9O6K) |
| **Data collection and processing** | |
| Microscope | Titan Krios |
| Camera | Gatan K3 |
| Magnification | 81,000 |
| Voltage (kV) | 300 |
| Defocus range (µm) | -0.4 ~ -3.1 |
| Exposure time (s) | 2 |
| Dose rate($e^{-}$/Å$^2$/s) | 26.77 |
| Number of frames | 50 |
| Pixel size (Å) | 0.529 |
| Micrographs (no.) | 5137 |
| Symmetry imposed | *C1* |
| Initial particles (no.) | 2,645,958 |
| Final particles (no.) | 66,922 |
| Map resolution (Å) | 3.59 |
| FSC threhold | 0.143 |
| **Refinement** | |
| Initial model used | AMT1, AMT7, AMTP2 (7YI8)<br>AMTP1 (AlphaFold) |
| Model resolution (Å) | 3.8 |
| FSC threshold | 0.143 |
| Map sharpening *B* factor (Å$^2$) | -100 |
| CC (mask) | 0.62 |
| Model composition | |
| Non-hydrogen atoms | 5847 |
| Protein residues | 756 |
| SAH | 1 |
| *B* factors (Å$^2$) | |
| Protein | 230.25 |
| SAH | 146.36 |
| R.m.s. deviations | |
| Bond lengths (Å) | 0.004 |
| Bond angles (°) | 0.775 |
| **Validation** | |
| MolProbity score | 1.95 |
| Clashscore | 9.69 |
| Poor rotamers (%) | 0.34 |
| Ramachandran plot | |
| Favored (%) | 93.22 |
| Allowed (%) | 6.78 |
| Disallowed (%) | 0.00 |
